# Redefining FLASH RT: the impact of mean dose rate and dose per pulse in the gastrointestinal tract

**DOI:** 10.1101/2024.04.19.590158

**Authors:** Kevin Liu, Trey Waldrop, Edgardo Aguilar, Nefetiti Mims, Denae Neill, Abagail Delahoussaye, Ziyi Li, David Swanson, Steven H. Lin, Albert C. Koong, Cullen M. Taniguchi, Billy W. Loo, Devarati Mitra, Emil Schüler

## Abstract

**Background:** The understanding of how varying radiation beam parameter settings affect the induction and magnitude of the FLASH effect remains limited.

**Purpose:** We sought to evaluate how the magnitude of radiation-induced gastrointestinal (GI) toxicity (RIGIT) depends on the interplay between mean dose rate (MDR) and dose per pulse (DPP).

**Methods:** C57BL/6J mice were subjected to total abdominal irradiation (11-14 Gy single fraction) under conventional irradiation (low DPP and low MDR, CONV) and various combinations of DPP and MDR up to ultra-high-dose-rate (UHDR) beam conditions. The effects of DPP were evaluated for DPPs of 1-6 Gy while the total dose and MDR were kept constant; the effects of MDR were evaluated for the range 0.3– 1440 Gy/s while the total dose and DPP were kept constant. RIGIT was quantified in non-tumor–bearing mice through the regenerating crypt assay and survival assessment. Tumor response was evaluated through tumor growth delay.

**Results:** Within each tested total dose using a constant MDR (>100 Gy/s), increasing DPP led to better sparing of regenerating crypts, with a more prominent effect seen at 12 and 14 Gy TAI. However, at fixed DPPs >4 Gy, similar sparing of crypts was demonstrated irrespective of MDR (from 0.3 to 1440 Gy/s). At a fixed high DPP of 4.7 Gy, survival was equivalently improved relative to CONV for all MDRs from 0.3 Gy/s to 104 Gy/s, but at a lower DPP of 0.93 Gy, increasing MDR produced a greater survival effect. We also confirmed that high DPP, regardless of MDR, produced the same magnitude of tumor growth delay relative to CONV using a clinically relevant melanoma mouse model.

**Conclusions:** This study demonstrates the strong influence that the beam parameter settings have on the magnitude of the FLASH effect. Both high DPP and UHDR appeared independently sufficient to produce FLASH sparing of GI toxicity, while isoeffective tumor response was maintained across all conditions.

## INTRODUCTION

Ultra-high-dose-rate (UHDR, ≥40 Gy/s) radiation therapy (RT) selectively spares normal tissue while maintaining isoeffective tumor killing *in vivo* (i.e., the FLASH effect) relative to conventional (CONV) dose-rate (CDR, <1 Gy/s) RT(1-5). Although the FLASH effect has been reported in different organ systems (e.g., brain, lung, gastrointestinal [GI] tract, skin) and animal species (e.g. rodents, human, swine, feline)(3,6-14), some studies have reported either no effect or increased detrimental effects on normal tissue from UHDR relative to CDR irradiation(15-19). Furthermore, even in reports showing increased normal tissue sparing after FLASH relative to CONV, the magnitude of the relative amount of sparing was inconsistent between studies and between institutions(4,6,17). Causes for these apparently conflicting results are not well understood; however, defining the FLASH effect based only on a single beam parameter (that is, a mean dose-rate [MDR] exceeding 40 Gy/s) may be an oversimplification. Rather, other aspects of the radiation beam including radiation source (x-rays, protons, electrons, heavy ions), dose per pulse (DPP), pulse repetition frequency (PRF), pulse width (PW), total delivery time, irradiation volume, and intrapulse dose rate (IDR; the dose rate within a pulse) may all contribute to the FLASH effect(4,20,21). Indeed, when matching beam parameters were used, the FLASH effect has been shown to be reproducible between institutions using the same biological assay(10,11).

To understand how individual radiation beam parameters are related to normal tissue sparing and tumor response, detailed dosimetric data are needed alongside the biological data. Many preclinical studies involving UHDR deliveries have failed to provide comprehensive descriptions of the irradiation parameters and setups, thereby limiting the prospects of retrospective analysis(4). The prevalence of such inconsistencies has contributed to what is now recognized as the ‘reproducibility crisis’ in radiobiological data within the field of radiation oncology (22-24). To safeguard the clinical translation of FLASH, the robustness and reproducibility of the FLASH effect need to be established.

To start addressing the question of how the interplay of various radiation beam parameters affects the induction and magnitude of the FLASH effect, we used a combinatorial approach to determine the influence of DPP and MDR on GI toxicity (in non-tumor–bearing mice) and on tumor response (in a clinically relevant tumor model system). An electron irradiator was used to allow full control of beam parameters and to isolate the effects of DPP and MDR. We demonstrated that radiation-induced gastrointestinal toxicity (RIGIT) is reduced compared to CONV by either high DPP or UHDR delivery, each apparently independently sufficient. We also confirmed the iso-efficacy of all the optimized beam conditions on tumor response.

## METHODS

### Mouse handling

Eight-week-old female C57BL/6J mice from Jackson Laboratories (Bar Harbor, Maine, USA) were used for these experiments. Mice were acclimatized for a minimum of 3 days before the experiment was begun. The mice had access to standard chow (PicoLab Rodent Diet 20, no. 5053; PMI Nutrition International, St. Louis, MO, United States) and acidified water ad libitum. Five mice were housed per ventilated cage in a 12/12-hour light/dark cycle. All mouse procedures used were approved by the Institutional Animal Care and Use Committee (IACUC) at our institution.

GI toxicity was evaluated with the crypt regeneration assay (n=5-10 mice/group) and with downstream survival analyses (n=10 mice/group). In the crypt assay, mice were euthanized at 84 hours after irradiation via CO_2_ inhalation followed by cervical dislocation. The jejunum from each mouse was collected, washed with phosphate-buffered saline, fixed in 10% neutral buffered formalin for 24 hours, and transferred to 70% ethanol after 24 hours of fixation. The jejunum was then sliced into nine 3-μm-thick transverse sections and stained with hematoxylin and eosin and embedded in paraffin. Regenerating crypts were counted manually in transverse sections that included complete jejunal circumferences. Regenerating crypts were defined as having (1) basophilic structure along the circumferential edge, (2) U-shaped structure, and (3) at least 10 cells(9,10).

An additional 10 mice per group were followed for up to 86 days after irradiation for survival analyses. Throughout this period, indicators of acute toxicity, such as weight loss, hunching, diarrhea, activity, and behavioral changes, were closely observed. Early euthanasia (before the completion of the experimental period) was implemented if mice met the following IACUC criteria: (1) displaying moribund characteristics (e.g., hunched posture, non-weight-bearing lameness, ruffled fur, labored breathing) or (2) experiencing a loss exceeding 30% of the baseline body weight. On the 86^th^ day, all remaining mice were euthanized to complete the experiment.

### Irradiation setup

All experiments were performed with an IntraOp FLASH Mobetron electron linear accelerator (IntraOp Medical, Sunnyvale, CA, USA) that delivered 9-MeV electrons. A custom-made mouse cradle was designed and 3D-printed to immobilize the mice while they were sedated with isoflurane in ambient air. Mouse positioning in the cradle was verified by CT imaging on a small animal radiation research platform, using a different cohort of age and sex matched mice (SARRP, Xstrahl Inc., Swanee, GA, USA). A custom-made collimator was produced and seated at the exit window of the Mobetron, as described elsewhere,(10,25) to allow sedation and reproducible setup placement for full and consistent total abdominal irradiation to a field size of 40 x 40 mm^2^ (**Fig. 1**). The irradiation setup was calibrated for each irradiation condition to a depth of 8 mm. Dosimetric calibration was done with dose-rate-independent Gafchromic EBT3 film paired with beam current transformers, the latter to allow real-time readout of the individual radiation beam parameters for each delivery (**Table S1**). During the FLASH irradiations, the beam current transformers were used to monitor the dose delivery and to log all radiation beam parameters on a per-pulse basis for each individual mouse(10,26-29). All CONV irradiations were done at a set MDR of 0.3 Gy/s and a DPP of 10 mGy.

**Fig. 1.**
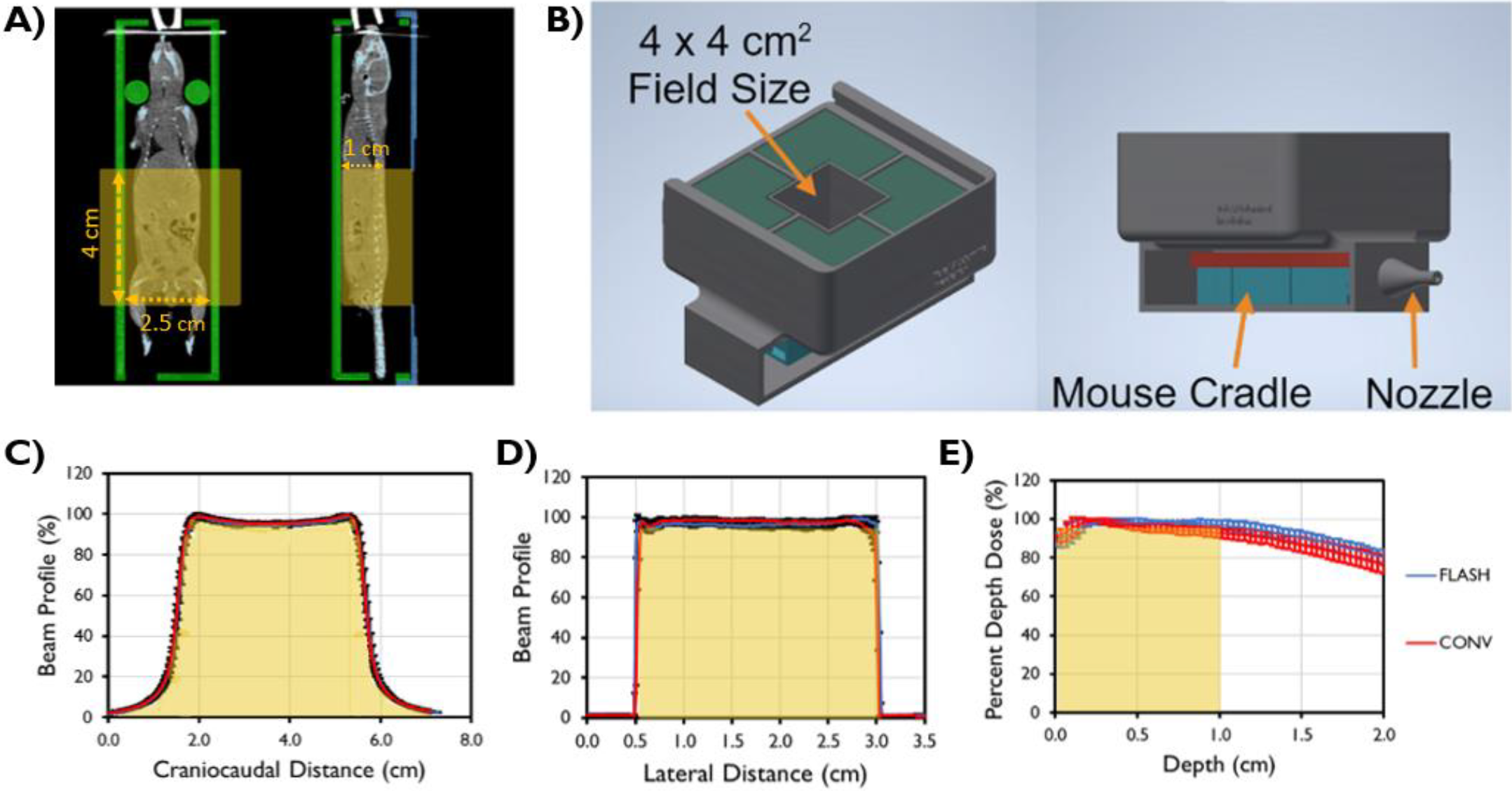
Experimental platform used for total abdominal irradiation. (A) Coronal and sagittal CT slices of a mouse placed in the mouse cradle, with columns used to support the neck for reproducible positioning. (B) The 3D platform used to collimate the field and index the mouse cradle to the radiation field. (C,D) Craniocaudal and lateral profiles at a depth of 8 mm in solid water placed inside the cradle, and (E) depth dose profiles for FLASH and CONV irradiation as measured with Gafchromic EBT3 film, demonstrating uniform dose throughout the abdominal treatment volume. Yellow shading indicates where the mouse was irradiated in the CT images and in the dose profiles.

### Evaluating irradiation parameters for acute GI toxicity

#### Evaluating dose per pulse at constant mean dose rate

The extent to which the magnitude of RIGIT depends on DPP was evaluated by delivering DPPs of 1–5.5 Gy (total dose 11 Gy), 1.3–6 Gy (total dose 12 Gy), and 1.08–4.7 Gy (total dose 14 Gy). The range of DPP values was based on the machine output and available settings to ensure a constant MDR within each dose settings. The DPP was modified by changing the PW setting while keeping the IDR constant. Variation in MDR within each dose group was minimized by adjusting the PRF for each delivery. The MDR (± SD) for the 11-Gy dose group was 161 ± 12 Gy/s; for the 12-Gy group, 184 ± 3 Gy/s; and for the 14-Gy group, 104 ± 2 Gy/s (**Table S1**).

#### Evaluating mean dose rate at constant dose per pulse

The dependence of the magnitude of RIGIT on MDR was evaluated by delivering 11 Gy, 12 Gy, or 14 Gy at corresponding constant DPPs of 5.5 Gy, 6 Gy, or 4.7 Gy, respectively. The MDR range investigated within each dose group was 0.3–1350 Gy/s for 11 Gy, 0.3–1440 Gy/s for 12 Gy, and 0.3–830 Gy/s for 14 Gy. The MDR was modified by adjusting the time between pulses by changing the PRF from 5 to 120 Hz. For MDRs that fell below the PRF setting of 5 Hz (e.g., ∼40 Gy/s, the lowest PRF setting on the Mobetron system), an in-house–developed control board was used to further increase the time between pulses while at the same time maintaining full control over all other radiation beam parameters, as done in a recent study (30).

### Evaluating radiation beam parameters for survival and tumor response

#### Survival of mice treated with varied dose per pulse and mean dose rate settings

For the survival analysis, mice were irradiated to a dose of 14 Gy under the following conditions (n=10/group): High-DPP (FLASH), Low-DPP (FLASH), High-DPP (CONV), Low-DPP (CONV), and standard CONV. For the High-DPP conditions, the DPP was 4.7 Gy; for the Low-DPP conditions, the DPP was 0.93 Gy. The MDR for the FLASH groups was 104 Gy/s and that for the CONV groups was 0.3 Gy/s. The PRF settings used for the High-DPP (FLASH) and Low-DPP (FLASH) groups were 15 and 105 Hz, respectively, to maintain a constant MDR of 104 Gy/s. The High-DPP (CONV) and Low-DPP (CONV) dose rate of 0.3 Gy/s was obtained by equidistant temporal delivery of the pulses into three groups. The parameters for standard CONV irradiations were 10 mGy and 0.3 Gy/s.

#### Irradiation of subcutaneous tumors

D4M murine melanoma cells were acquired (31). This stable cell line was derived from a single-cell clone of the D4M.3A melanoma cell line, established from a *Tyr:CreER;Braf*^*CA*^*;Pten*^*lox/lox*^ mouse.(32,33) Cells were incubated at 5% CO_2_ at 37°C and cultured in DMEM-high glucose medium (Sigma-Aldrich, St. Louis, MO) with 10% fetal bovine serum and 1% penicillin and streptomycin. Cells were confirmed as negative for mycoplasma contamination by using a Lonza Mycoplasma Detection Kit (Lonza Bioscience, Durham, NC) before injection. Aliquots of 10^6^ cells/100 µL were prepared and injected subcutaneously into the flank of the mice under isoflurane anesthesia. When tumor volumes reached roughly 100 mm^3^, mice were randomly assigned to a 0-Gy SHAM group or a 20-Gy treatment group under the following DPP and MDR conditions: CONV (10 mGy, 0.3 Gy/s); High-DPP FLASH (6.7 Gy, 1208 Gy/s); or High-DPP CONV (6.7 Gy, 0.3 Gy/s), with n = 5-6 mice per group. Starting on day 0 (day of irradiation), tumor size was monitored 2-3 times per week with calipers for 30 days or until euthanasia. Tumor growth time and tumor growth delay were assessed by measuring and plotting the time until each individual tumor volume reached 2, 3, and 5 times its initial volume at the time of irradiation.

### Statistical analyses

To assess normal tissue toxicity, the number of regenerating crypts from the FLASH- and CONV-treated groups were compared by two-sample using t tests. A linear least squares linear regression model was used to identify trends in crypt count as a function of DPP or MDR. To assess tumor growth effects, the tumor growth time and tumor growth delay between treatment groups were compared with Kruskal-Wallis H tests. All statistical analyses were done with GraphPad Prism V.10 (La Jolla, CA, USA) and R V4.3.2(Auckland, NZ), with P values of <0.05 considered to indicate significant differences. Benjamini & Hochberg adjustment has been used to control for multiple testing.

## RESULTS

The measured radiation beam parameters delivered in this study are summarized in Table S1. Acute GI toxicity, evaluated with a regenerating crypt assay, decreased with increasing DPP across all dose groups while the MDR was held constant between different DPPs within the same dose setting (**Fig. 2**). At 12 Gy, crypt counts after FLASH were significantly higher than after CONV for all evaluated DPPs (from 1.3 to 6 Gy). At 14 Gy, crypt counts after FLASH were significantly higher at the higher DPPs (from 2 to 4.7 Gy). In comparing the relative difference in response between the highest DPP groups for the 11 Gy (DPP=5.5 Gy), 12 Gy (DPP=6 Gy), and 14 Gy (DPP=4.7 Gy) dose groups, we found that the mean numbers of regenerating crypts counted were 1.5 times, 2 times, and 2.9 times higher for FLASH versus CONV. At total doses of 12 Gy and 14 Gy, the mean number of regenerating crypts per circumference increased linearly with DPP (*P*=0.0024 for 12 Gy and *P*=4.13 e-05 for 14 Gy).

**Fig. 2.**
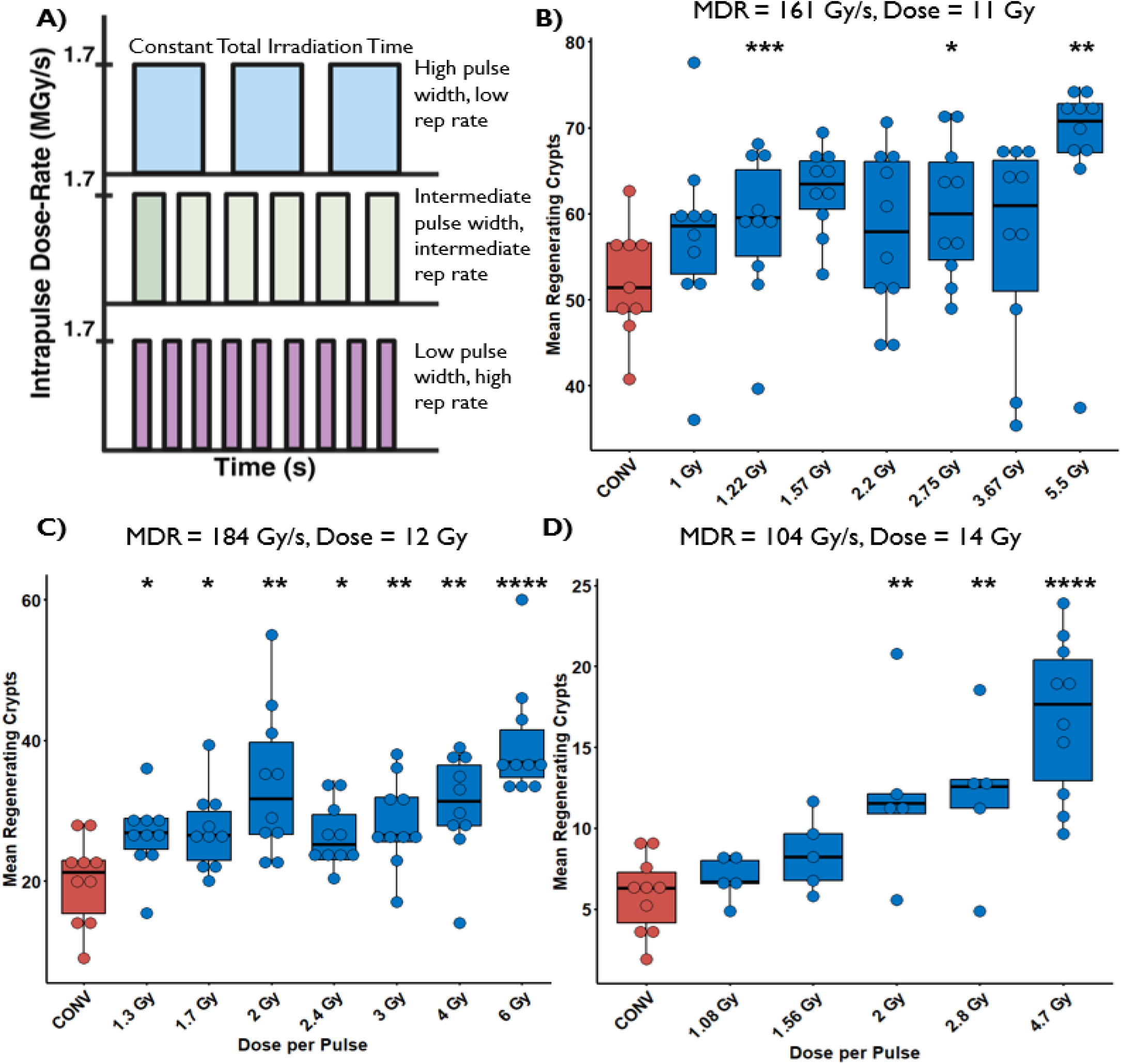
(A) The beam structure schema (not to scale) illustrating how pulse width (PW) and pulse repetition frequency (PRF) are varied to achieve a delivery of different dose per pulse (DPP) while maintaining constant intrapulse dose rate, total irradiation time, mean dose rate (MDR), and total dose for each set of experiments in B-D. The mean number of regenerating crypts per circumference for mice receiving a total abdominal irradiation by either CONV - (0.3Gy/s, 10mGy/pulse, Red) or FLASH (Blue) to a dose of B) 11 Gy, MDR = 161 Gy/s, C) 12 Gy, MDR = 184 Gy/s, and D) 14 Gy, MDR = 104 Gy/s. An unpaired parametric t-test was used to compare the FLASH treated groups with the CONV treated group.^*^ p< 0.05; ^**^ p < 0.01; ^***^ p < 0.001; ^****^ p < 0.0001. For doses of 11-14 Gy and MDR of 104-184 Gy/s, sparing of crypts increases progressively with increasing DPP, especially at 14 Gy/104 Gy/s. At the lower doses/higher MDRs of 11-12Gy/161-184 Gy/s tested here; significant sparing of crypts is observed at DPPs > 1 Gy.

### The magnitude of normal tissue sparing in high DPP conditions is independent of the time to deliver a given dose (mean dose rate)

When the highest DPP setting available was used to deliver a total abdominal dose of 11–14 Gy, the magnitude of RIGIT was unaffected by the amount of time between pulses and thereby independent of the MDR and total irradiation time (**Fig. 3**). The crypt count in high-DPP FLASH-treated mice was significantly higher than in CONV-treated mice for all MDR groups across all doses delivered. When comparing the mean number of regenerating crypts under a given DPP setting with varied MDRs, we found that the MDR did not influence the number of crypts in the high-DPP beam conditions, even at 0.3 Gy/s. Under high DPP conditions, no statistically significant association was found between MDR and the number of regenerating crypts at any of the dose levels tested.

**Fig. 3.**
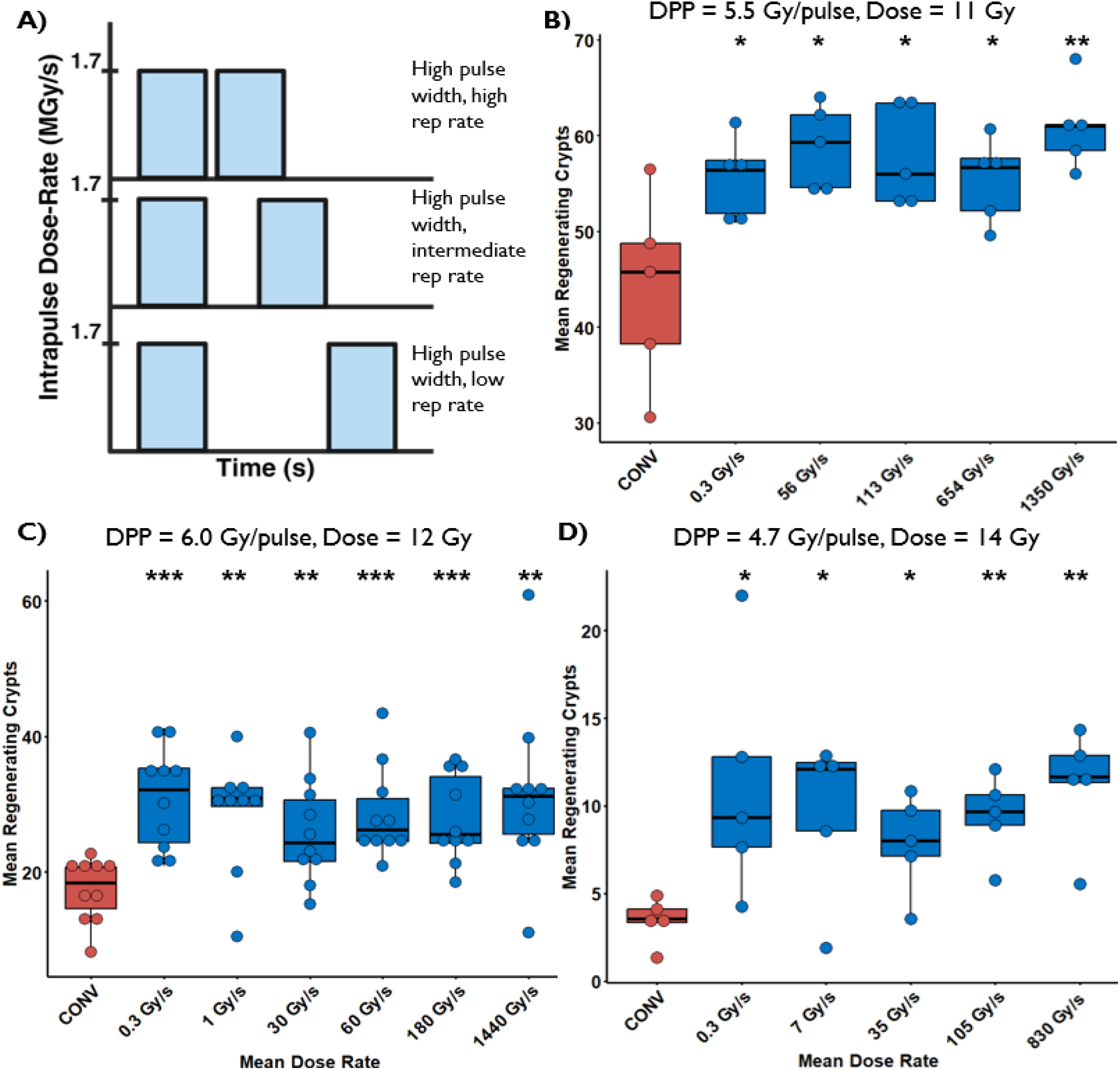
(A) The beam structure schema (not to scale) illustrating how the interval between two pulses of constant dose per pulse (DPP), pulse width and intrapulse dose rate is varied to produce varying total irradiation time and mean dose rate (MDR) for each set of experiments in B-D. The mean number of regenerating crypts per circumference for mice receiving a total abdominal irradiation by either CONV (0.3 Gy/s, 10 mGy/pulse) or high DPP irradiation to a dose of B) 11 Gy, DPP = 5.5 Gy per pulse C) 12 Gy, DPP = 6 Gy per pulse, and D) 14 Gy, DPP = 4.7 Gy per pulse. An unpaired parametric t-test was used to compare the high DPP treated groups with the CONV treated group. ^*^ p< 0.05; ^**^ p < 0.01; ^***^ p < 0.001; ^****^ p < 0.0001. For DPP of 4.7-6 Gy, sparing of crypts compared to CONV at doses of 11-14 Gy was observed for all MDR from 0.3-1440 Gy/s.

### Survival after FLASH is influenced by both the dose per pulse and mean dose rate

The pulse structure of the High-DPP (FLASH) group involved temporally equidistant delivery of three pulses (4.7 Gy per pulse) over a period of 0.13 seconds, whereas the High-DPP (CONV) group involved the equidistant delivery of three pulses (4.7 Gy per pulse) over a period of 45 seconds for CDR (**Fig. 4A,B**). The pulse structure of the Low-DPP (FLASH) group involved equidistant delivery of 15 pulses (0.93 Gy per pulse) over a period of 0.13 seconds, and the Low-DPP (CONV) group involved the equidistant delivery of three sets of five pulses (4.7 Gy delivered per set, 0.93 Gy per pulse) over a period of 45 seconds (**Fig. 4A,B**). The High-DPP and Low-DPP FLASH groups lost weight at a slower rate than the groups treated at CDR regardless of DPP; however, no significant difference in total weight lost was noted between groups (**Fig. 4C**). The standard CONV was delivered at a DPP of 10 mGy and an MDR of 0.3 Gy/s. At 86 days after irradiation, mice receiving high-DPP irradiation (4.7 Gy/pulse) had similar survival rates (50%-60%) regardless of whether the radiation was delivered under UHDR or CDR conditions. The survival rate for the Low-DPP (0.93 Gy/pulse) group treated with UHDR was also 60%. However, mice treated with Low-DPP at CDR had lower survival (10%-20%), as did those treated with standard CDR (10 mGy DPP, 0.3 Gy/s).

**Fig. 4.**
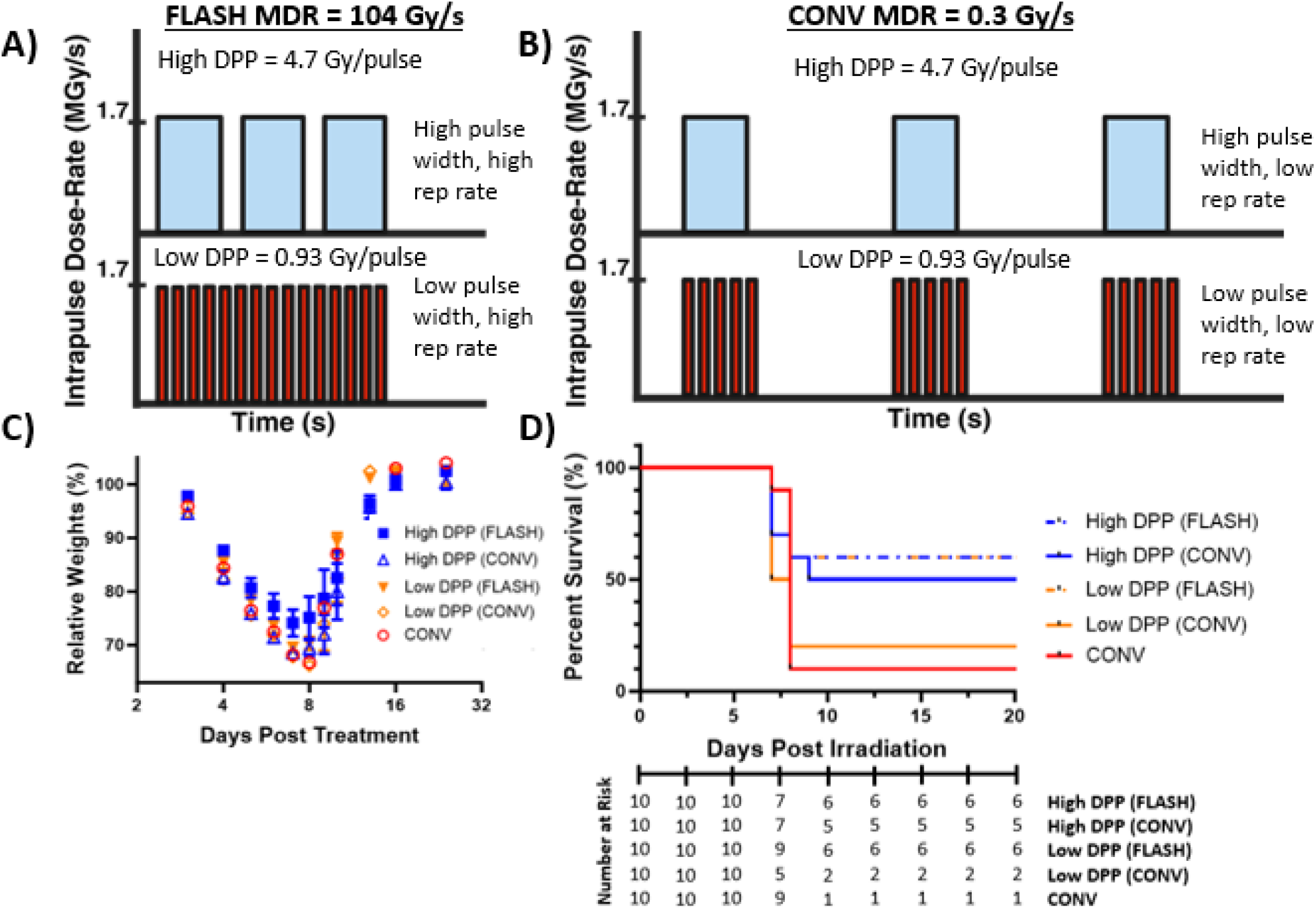
The beam structure schema (not to scale) showing constant dose delivery for a High DPP (4.7 Gy) and Low DPP (0.93 Gy) beam at (A) FLASH dose-rates and (B) CONV dose-rates. (C) Plot of the relative weights at up to 24 days post irradiation for the treated groups. (D) Kaplan–Meier plots shows percent survival paired with the Number at Risk as a function of days post treatment with FLASH or CONV after 14 Gy total abdominal irradiation (n = 10/group). The high DPP FLASH and CONV delivery consisted of 3 pulses delivered with a pulse repletion frequency (PRF) of 15 and 0.03, respectively. The low DPP FLASH irradiation was delivered with 15 pulses (PRF = 105 Hz) while the low DPP CONV irradiation was delivered with 3 sets of 5 pulses, each set delivered with a PRF of 0.233 Hz. Increased survival compared to CONV was observed with both high DPP deliveries (including FLASH and CONV) as well as with both FLASH deliveries (high and low DPP), but not low DPP (CONV).

### Isoeffective tumor response is maintained across all irradiation conditions

To evaluate the effects of altering beam parameters on tumor growth kinetics, mice bearing subcutaneous D4M melanoma tumors were irradiated with 20 Gy and followed for tumor growth (**Fig. 5**). All irradiation parameters tested significantly increased the time to reach 2x, 3x and 5x pre-irradiation volume, relative to sham-treated mice. The one exception was an apparent trend (*P*=0.054) to delay in reaching 5x volume after standard CONV treatment (0.01 Gy/pulse, 0.3 Gy/s). However, no significant differences in time to reach these volume thresholds were found between the three beam parameters of standard CONV, High-DPP FLASH (6.7 Gy/pulse, 1208 Gy/s), and High-DPP CONV (6.7 Gy/pulse, 0.3 Gy/s). Consistent with these results, no significant differences in tumor growth delay were found between irradiated tumors regardless of beam parameters, with a constant offset of 4-5 days between the mean amount of time to reach a tumor volume relative to the mean amount of time in the sham group.

**Fig. 5.**
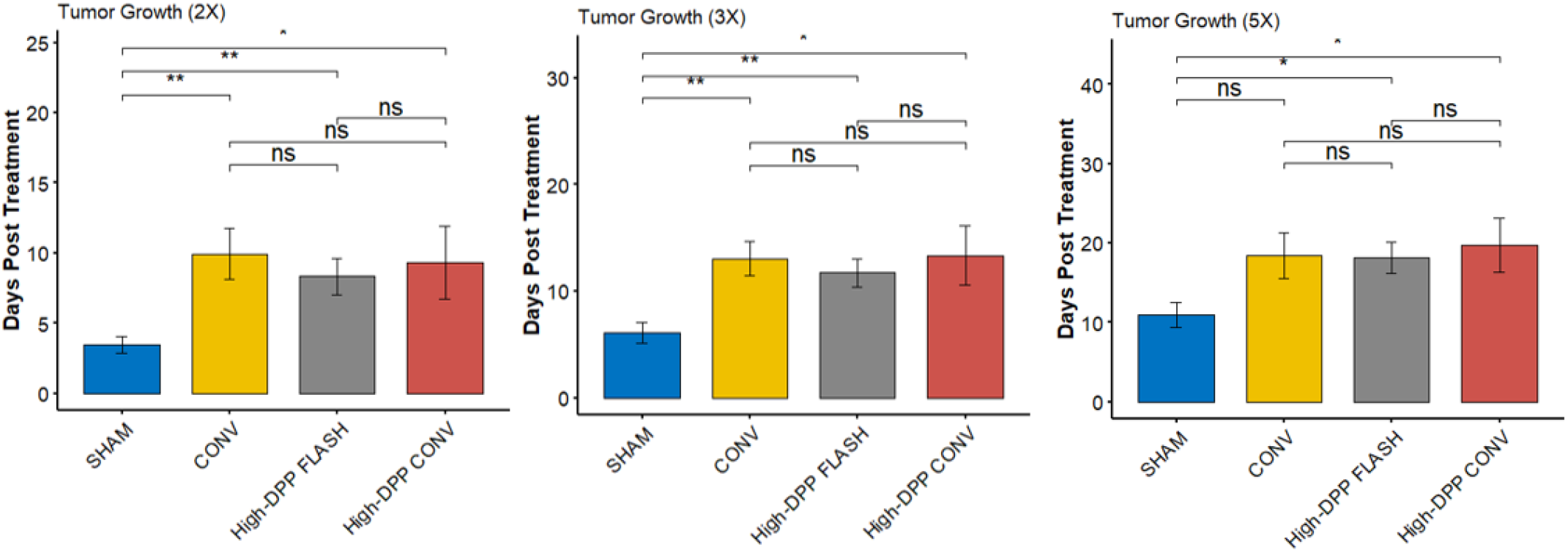
Time for subcutaneous D4M tumors to reach 2-5 times their original volume after focal tumor irradiation to a dose of 20 Gy under conventional (CONV), High-dose per pulse (DPP) CONV, or High-DPP FLASH conditions. The MDR values were 0.3 Gy/s for CONV, 0.3 Gy/s for High-DPP CONV, and 1208 Gy/s for High-DPP FLASH; the corresponding DPPs were 0.01 Gy, 6.7 Gy, and 6.7 Gy. Error bars represent standard error of the mean. FLASH-treated vs CONV-treated groups were compared for tumor growth time with unpaired parametric *t* tests, and for tumor growth delay with Kruskal-Wallis tests. ^*^*P*<0.05; ^**^*P*<0.01; ^***^*P*<0.001; ^****^*P*<0.0001. No significant differences on tumor growth delay were found between the treated groups.

## DISCUSSION

The current study is the first, to our knowledge, to demonstrate that high dose per pulse (DPP) and ultra-high dose rate (UHDR) conditions independently impact the magnitude of normal tissue toxicity with FLASH irradiation. The total dose delivered to the abdomen in this study was selected based on previous findings that 11 Gy produced similar toxicity between standard FLASH conditions (1-2 Gy/pulse, 100-200 Gy/s) and CONV (low DPP and low mean dose rate); that 12 Gy was the “middle ground” for a significant observable difference; and that 14 Gy was a lethal dose in which the regenerating crypt number in the conventional dose rate (CDR) group typically fell below 5 per jejunal circumference(10). The absence of difference in GI sparing between CONV and FLASH at 11 Gy was reproduced when similar DPPs were used(10). However, differential responses were found at higher DPP settings at this lower dose level. At doses of 12 and 14 Gy, we found the magnitude of sparing under UHDR conditions to be more pronounced at a wider range of DPP settings (1.3–6 Gy and 2–4.7 Gy per pulse, respectively). Differences in the mean dose rate (MDR) between different doses within the DPP evaluation in the current study were caused by constraints in the Mobetron system, given that the pulse repetition frequency (PRF) settings were fixed; however, our aim was to use FLASH dose rates in excess of 100 Gy/s, similar to those in other preclinical studies(7,10,12,30,34). We further found that larger numbers of crypts were present at the highest DPP setting for all doses investigated, with a positive statistical correlation with DPP indicating higher DPP yields greater sparing than lower DPP settings, even though MDR is kept constant. Moreover, when the highest DPP setting was maintained but the time between pulses was varied (thereby modifying the MDR and total irradiation time) no differences were found in the number of regenerating crypts, even at dose rates as low as 0.3 Gy/s, matching the CDR setting on the IntraOp Mobetron.

A recent summary of the temporal dosimetry characteristics of FLASH in *in vivo* and *in vitro* experiments with different organs and biological endpoints(6) indicated that a combination of higher IDRs (>10^5^ Gy/s) and shorter irradiation times (<1 s) is necessary to evoke a FLASH-sparing effect. In addition to the IDR and irradiation time, another series of experiments revealed that MDR was also important for neuroprotection after whole-brain FLASH irradiation to 10 Gy, with the threshold for such an effect being 30 Gy/s(7). Another group found an increase in toxicity sparing as a function of MDR after total abdominal irradiation(35); in that study, the dose lethal to 50% of the study population (LD_50_) was 14.7 Gy when the MDR was 0.5 Gy/s, 16.6 for an MDR of 70 Gy/s, and 18.3 Gy for an MDR of 210 Gy/s. However, other studies have failed to see a FLASH sparing effect, or in some cases have shown even greater toxicity, under similar MDR conditions(15,17,36-38), clearly demonstrating the inadequacy of defining FLASH based solely on MDR.

Our finding that high DPP produced similar toxicity regardless of MDR and total irradiation time (Fig. 3) contradicts findings from previous studies, such as one that reports an MDR threshold of 280 Gy/s for significantly more sparing of regenerating crypts compared to the CONV-treated group when the total dose to the abdomen was 11.2 Gy delivered with a DPP of 5.6 Gy per pulse.(39) This study also reported significantly more regenerating crypts at a lower DPP setting of 2.24 Gy (vs 5.6 Gy) when the MDR was kept constant at 280 Gy/s. The inconsistencies between our findings despite using similar beam parameters for the regenerating crypt assay will require further study but may reflect the use of different types of mice (C3H vs C57/BL6), especially in the gut microbiome(40), or differences in beam energy (6 MeV vs 9 MeV) and how the output and temporal structure of the beam was modified despite having IDRs, DPPs, and MDRs compatible with those in this study(4). In the prior study, a 6-MeV beam was used with the output and dose rate of the beam varied by adjusting the scattering foil-to-collimator distance and lowering the electron gun current while keeping a constant PW, thus changing the IDR (ranging from 2.2-5.9 MGy/s). In our study, the output and dose rate were modified by adjusting the pulse width (PW) and PRF settings and keeping the IDR (1.5-1.7 MGy/s) constant, suggesting that future investigations into how the beam parameter settings influence the FLASH effect should include the IDR. Additionally, there was also a difference in how the regenerating crypts were quantified. The prior study only scored the crypts in a segment of the intestine that had the most amount of damage (least amount of crypts), while in the current study the number of crypts were counted and averaged along nine different 3-mm sections of the jejunum. These discrepancies emphasize the need for future investigations into the reproducibility of the FLASH effect between different types of linear accelerators between institutions under identical beam conditions and identical biological assays.

Our evaluations of mouse survival after total abdominal irradiation with different DPP and MDR combinations revealed no differences in survival after a DPP of 4.7 Gy per pulse but at different mean dose-rates (104 Gy/s vs 0.3 Gy/s). However, at lower DPP settings (0.93 Gy per pulse), the MDR was found to affect survival, as a higher proportion survived after treatment at 104 Gy/s than at 0.3 Gy/s. Mice treated at identical MDRs (104 Gy/s) but at different DPPs (4.7 Gy vs 0.93 Gy) also had no difference in survival, suggesting that normal-tissue sparing depends on MDR at low DPP settings, whereas at higher DPP settings the MDR dependence diminishes, indicating that high DPP or UHDR irradiations are independently sufficient to produce the FLASH sparing effect.

To examine if a high DPP beam setting at UHDR or CDR yields isoeffective tumor response, we used a subcutaneous melanoma mouse model (D4M) to examine the time to tumor doubling, tripling, and quintupling. Unlike several other commonly used melanoma mouse models, the D4M model carries two common genetic drivers of human melanoma and as a result may be particularly clinically relevant(32). This is the first study, to our knowledge, to examine the response of D4M tumors in a FLASH beamline. In evaluating the efficacy of FLASH vs CONV on tumor control in this model, we found equivalent tumor responses after CONV, High-DPP FLASH, and High-DPP CONV treatment to the same total dose, despite treatment at DPPs ranging from 0.01-6.7 Gy/pulse and MDRs ranging from 0.3-1208 Gy/s. This finding is particularly relevant for FLASH, as many papers have shown FLASH to be as effective, or even more effective, than CONV against a variety of tumor types. Iso-efficacy has been demonstrated in several tumor models, including lung adenocarcinoma(40), glioblastoma(6), and ovarian carcinoma(21, 41). This systematic analysis examined datasets from 15 publications describing various tumor types, irradiation modalities, and endpoints and concluded that FLASH was, on average, as effective as CONV treatments. Those authors also noted no correlation between beam parameters(42); however, they also examined fewer parameters than did the current study.

It is important to note that this finding highlighting the unique effects of DPP and MDR was only investigated for RIGIT and that such effects must be confirmed in other organ systems using other biological assays. A study investigating the effects of MDR on whole brain irradiation of mice, in which an electron beam (16 MeV) with a DPP of 1.75 Gy delivered at UHDR (200-300 Gy/s) and CDR (0.13 Gy/s) was used, found that irradiations at UHDR with a DPP of 1.75 Gy yielded reduced cognitive deficits compared to those with CDR deliveries despite identical DPP settings(41). This may indicate that for whole brain irradiations that a DPP of 1.75 Gy may be insufficient to bring about a significant sparing effect independent of the MDR. However, in a different study that involved whole brain irradiation with FLASH and CONV treatments, nonsignificant differences were found in the recognition ratio of mice treated with a high DPP FLASH (10 Gy/pulse, 5.5E6 Gy/s) and low DPP FLASH (1 Gy/pulse, 110 Gy/s). Both groups showed significant differences when compared against mice treated with CONV (0.1 Gy/s), which is comparable to our findings on the survival of mice treated with a high DPP FLASH (4.7 Gy/pulse, 104 Gy/s) and low DPP FLASH (0.93 Gy/pulse, 104 Gy/s) for abdominal irradiations(42). Additionally, the prior study also found preferential sparing in mice treated with continuous delivery of proton FLASH (110 Gy/s) compared to proton CONV (0.1 Gy/s), which has also been reported for other tissues and endpoints(12,13,43). Based on these prior data and the data presented here, the correlation between the chosen radiation beam parameters and the magnitude of the FLASH effect is clear, but the exact parameter settings and magnitude of sparing is not. Based on the data presented here, for low DPP settings (< 4 Gy), a FLASH effect can only be induced at sufficiently high MDRs (typically 100 Gy/s as demonstrated in prior publications) while a high DPP setting (≥ 4 Gy) can induce a FLASH sparing effect regardless of the MDR used, which could help inform on the clinical use and translation of FLASH. Conformal CONV, using approaches other than electron irradiation, generally involves the delivery of radiation from many different angles to maximize the dose delivered to the tumor while minimizing the dose to the patient’s normal tissue. Doing so involves the slow rotation of a treatment gantry. Delivering a conformal treatment under FLASH conditions using a traditional delivery strategy may be infeasible given the slow rotation of standard gantries. As a result, the dose-rate of such standard treatments would be drastically reduced unless the beam was delivered in a multidirectional fashion simultaneously as proposed by the PHASER system(44). Achieving a clinical plan with high DPP could mitigate the issue of gantry movement, since the sparing effect would no longer be dependent on MDR. However, there remain significant uncertainties regarding how to practically deliver conformal FLASH, including issues of feasibility and confirmation that splitting the dose delivery does not reduce or eliminate the sparing effect, which has been suggested by others(45).

## CONCLUSIONS

Here we investigated how the choice of MDR and DPP in UHDR irradiations affects the magnitude of radiation-induced toxicity and tumor response. We observed that increasing the DPP, at a constant MDR, was linearly correlated with a reduction in toxicity across various dose levels, highlighting the importance of DPP in modulating the FLASH effect. We also found that using a DPP of >4 Gy led to a reduction in toxicity (vs CONV) regardless of the amount of time to deliver a given dose (MDR), which was not found at lower DPPs. Isoeffective tumor response using a clinically relevant murine melanoma model was maintained across optimized beam parameter conditions. These findings suggests that both high DPP and ultra-high MDR conditions appear to be independently sufficient to achieve the FLASH sparing effect. Overall, our study underscores the need for comprehensive reporting of dosimetric and biological data in studies involving UHDR radiation deliveries to enhance the reproducibility and translatability of FLASH to clinical settings. Further research is also needed to elucidate the specific contributions of different radiation beam parameters and their optimal combinations for maximizing the therapeutic benefits of FLASH while minimizing normal-tissue toxicity.

## Acknowledgements

We thank Christine F. Wogan, MS, ELS, of MD Anderson’s Division of Radiation Oncology, for editorial contributions to this article. We thank David Fisher, MD, PhD, of Massachusetts General Hospital’s Cutaneous Biology Research Center, and Michael Davies, MD, PhD, of MD Anderson’s Division of Cancer Medicine, for providing the D4M murine melanoma cells.

