## Supplemental Table 1 for "Redefining FLASH RT: the impact of mean dose rate and dose per pulse in the gastrointestinal tract"

***Supplementary Table S1:*** *Physical Beam Parameters Measured using Gafchromic Film and Beam Current Transformers*

| **Study** | **Dose(Gy)** | **Dose per Pulse (Gy)** | | **Mean Dose Rate (Gy/s)** | **Irradiation Time (s)** | **Instantaneous Dose Rate (MGy/s)** | **Pulse Width (µs)** | **Pulse Frequency (Hz)** | **Number of Pulses** |
| --- | --- | --- | --- | --- | --- | --- | --- | --- | --- |
| Dose per Pulse | 11.1 ± 0.1 | | 1.0 ± 0.0 | 133 ± 1 | 0.08 | 1.5 | 0.6 | 120 | 11 |
|  |  |  | 1.2 ± 0.0 | 166 ± 1 | 0.07 | 1.6 | 0.8 | 120 | 9 |
|  |  |  | 1.6 ± 0.0 | 167 ± 1 | 0.07 | 1.6 | 1.0 | 90 | 7 |
|  |  |  | 2.2 ± 0.0 | 168 ± 2 | 0.07 | 1.6 | 1.4 | 60 | 5 |
|  |  |  | 2.7 ± 0.0 | 164 ± 1 | 0.07 | 1.6 | 1.7 | 45 | 4 |
|  |  |  | 3.7 ± 0.0 | 166 ± 1 | 0.07 | 1.6 | 2.3 | 30 | 3 |
|  |  |  | 5.5 ± 0.0 | 166 ± 1 | 0.07 | 1.6 | 3.4 | 15 | 2 |
|  |  |  | 0.01 | 0.3 | 34 | 0.01 | 1.2 | 30 | 1025 |
| Mean Dose Rate | 11.2 ± 0.1 | | 5.6 ± 0.0 | 0.3 ± 0.0 | 34 | 1.7 | 3.3 | 0.03 | 2 |
|  |  |  | 5.6 ± 0.0 | 56 ± 0 | 0.20 | 1.7 | 3.3 | 5 | 2 |
|  |  |  | 5.7 ± 0.0 | 113 ± 1 | 0.10 | 1.7 | 3.3 | 10 | 2 |
|  |  |  | 5.6 ± 0.0 | 654 ± 4 | 0.02 | 1.6 | 3.3 | 30 | 2 |
|  |  |  | 5.5 ± 0.0 | 1349 ± 10 | 0.01 | 1.7 | 3.3 | 120 | 2 |
|  |  |  | 0.01 | 0.3 | 34 | 0.01 | 1.2 | 30 | 1025 |
| Dose per Pulse | 12.3 ± 0.2 | | 1.4 ± 0.0 | 187 ± 0 | 0.07 | 1.6 | 0.8 | 120 | 9 |
|  |  |  | 1.7 ± 0.0 | 183 ± 1 | 0.07 | 1.6 | 1.0 | 90 | 7 |
|  |  |  | 2.1 ± 0.0 | 187 ± 1 | 0.07 | 1.7 | 1.2 | 75 | 6 |
|  |  |  | 2.5 ± 0.0 | 185 ± 2 | 0.07 | 1.7 | 1.4 | 60 | 5 |
|  |  |  | 3.0 ± 0.0 | 182 ± 1 | 0.07 | 1.7 | 1.8 | 45 | 4 |
|  |  |  | 4.0 ± 0.1 | 180 ± 6 | 0.07 | 1.7 | 2.3 | 30 | 3 |
|  |  |  | 6.1 ± 0.0 | 182 ± 1 | 0.07 | 1.7 | 3.6 | 15 | 2 |
|  |  |  | 0.01 | 0.3 | 37 | 0.01 | 1.2 | 30 | 1107 |
| Mean Dose Rate | 12.0 ± 0.2 | | 6.0 ± 0.1 | 0.3 ± 0.0 | 37 | 1.7 | 3.6 | 0.03 | 2 |
|  |  |  | 6.0 ± 0.1 | 1 ± 0 | 12 | 1.7 | 3.6 | 0.08 | 2 |
|  |  |  | 6.0 ± 0.1 | 10 ± 0 | 1.2 | 1.7 | 3.6 | 0.83 | 2 |
|  |  |  | 6.0 ± 0.1 | 30 ± 0 | 0.4 | 1.7 | 3.6 | 2.50 | 2 |
|  |  |  | 6.0 ± 0.1 | 60 ± 1 | 0.2 | 1.7 | 3.6 | 5 | 2 |
|  |  |  | 6.0 ± 0.1 | 180 ± 3 | 0.07 | 1.7 | 3.6 | 15 | 2 |
|  |  |  | 5.9 ± 0.4 | 1440 ± 26 | 0.01 | 1.7 | 3.6 | 120 | 2 |
|  |  |  | 0.01 | 0.3 | 37 | 0.01 | 1.2 | 30 | 1138 |
| Dose per Pulse | 13.8 ± 0.3 | | 1.1 ± 0.0 | 106 ± 0 | 0.13 | 1.7 | 0.7 | 90 | 13 |
|  |  |  | 1.5 ± 0.0 | 104 ± 2 | 0.13 | 1.6 | 1.0 | 60 | 9 |
|  |  |  | 2.0 ± 0.0 | 103 ± 2 | 0.13 | 1.5 | 1.3 | 45 | 7 |
|  |  |  | 2.8 ± 0.1 | 103 ± 2 | 0.13 | 1.5 | 1.9 | 30 | 5 |
|  |  |  | 4.6 ± 0.1 | 103 ± 2 | 0.13 | 1.5 | 3.4 | 15 | 3 |
|  |  |  | 0.01 | 0.3 | 43 | 0.01 | 1.2 | 30 | 1280 |
| Mean Dose Rate | 14.0 ± 0.1 | | 4.6 ± 0.0 | 0.3 ± 0.0 | 45 | 1.5 | 3.1 | 0.04 | 3 |
|  |  |  | 4.7 ± 0.0 | 7 ± 0 | 2.0 | 1.5 | 3.1 | 1 | 3 |
|  |  |  | 4.7 ± 0.0 | 35 ± 0 | 0.4 | 1.5 | 3.1 | 5 | 3 |
|  |  |  | 4.7 ± 0.0 | 105 ± 1 | 0.13 | 1.5 | 3.1 | 15 | 3 |
|  |  |  | 4.7 ± 0.0 | 830 ± 4 | 0.02 | 1.5 | 3.1 | 120 | 3 |
|  |  |  | 0.01 | 0.3 | 45 | 0.01 | 1.2 | 30 | 1335 |
| Survival | 13.9 ± 0.2 | | 4.6 ± 0.0 | 104 ± 1 | 0.13 | 1.4 | 3.2 | 15 | 3 |
|  |  |  | 4.6 ± 0.0 | 0.3 ± 0.0 | 45 | 1.4 | 3.2 | 0.04 | 3 |
|  |  |  | 0.9 ± 0.0 | 104 ± 2 | 0.13 | 1.5 | 0.6 | 105 | 15 |
|  |  |  | 0.9 ± 0.0 | 0.3 ± 0.0 | 45 | 1.6 | 0.6 | 0.3 | 15 |
|  |  |  | 0.01 | 0.3 | 45 | 0.0 | 1.2 | 30 | 1356 |
| Tumor | 20.2 ± 0.1 | | 6.7 ± 0.0 | 1208 ± 6 | 0.02 | 1.7 | 3.96 | 120 | 3 |
|  |  |  | 6.7 ± 0.0 | 0.3 ± 0.0 | 60 | 1.7 | 3.96 | 0.03 | 3 |
|  |  |  | 0.01 | 0.3 | 60 | 0.01 | 1.2 | 30 | 808 |
